# Revisiting the SSRI–TRKB Mechanism: Lack of Evidence for Interaction in the Human Brain

**DOI:** 10.64898/2026.01.07.698082

**Authors:** Marie Klett, Sebastian Illes

## Abstract

Direct binding of selective serotonin reuptake inhibitors (SSRIs) to the TRKB receptor has been revealed as a novel mode of action for SSRIs. However, micromolar SSRI concentrations are required for this interaction to produce measurable effects in vitro and in vivo in animal models. Here, we highlight that measurements of human blood and cerebrospinal fluid from chronically treated patients reveal only nano- to picomolar SSRI levels. Rodent electrophysiological studies show that micromolar SSRI concentrations disrupt multiple ion channel functions, and we demonstrate that exposure of human neurons to micromolar concentrations leads to significantly reduced activity at 1 µM and complete silencing at 10 µM-concentrations required to exert effects by direct SSRI low-affinity binding to TRKB in animals.

Together, clinical measurements, animal electrophysiological studies, and our human neuro-electrophysiology data question whether the low SSRI concentrations achieved in patients have any consequences through their low-affinity binding to TRKB receptors. The micromolar concentrations required for effects elicit by SSRI-TRKB direct interaction are not reached under therapeutic conditions and, as our data show, would instead presumably lead to a near-complete loss of neuronal activity in humans.

## The SSRI-TRKB interaction is a novel mechanism that Operates Only at micromolar Concentrations of SSRIs

Selective serotonin reuptake inhibitors (SSRIs) are widely used to treat mood disorders. These agents are concentration-dependent polypharmacological drugs that, in addition to modulating SERT function, interact with several other molecular targets (Hyttel, 1993; Fagiolini, Cuomo and McIntyre, 2024; Li, 2024). Recent evidence suggests that SSRIs can bind to the TRKB receptor, promoting its translocation and facilitating BDNF dimerization, thereby contributing to neuroplasticity and behavioral changes observed in rodent models (Brunello, Cannarozzo and Castrén, 2024). This mechanism, however, requires micromolar SSRI concentrations. Specifically, fluoxetine (an SSRI) directly binds TRKB with micromolar affinity, and this binding is enhanced - but not competed - by cholesterol, suggesting a cooperative, dose-dependent interaction between SSRIs and cholesterol in TRKB recognition. Multiple antidepressants, including SSRIs, displace biotinylated fluoxetine from TRKB with micromolar *Ki* values, confirming direct, concentration-dependent binding. The TRKB.Y433F mutation preserves BDNF binding but impairs BDNF-induced TRKB Y816 phosphorylation, dimerization, lipid raft translocation, and FYN interaction, identifying a motif essential for SSRI-facilitated TRKB signaling at micromolar doses (Casarotto *et al.*, 2021; Casarotto, Umemori and Castrén, 2022). Overall, SSRIs at micromolar concentrations bind TRKB, and cholesterol levels modulate this interaction to facilitate TRKB surface localization and BDNF signaling in a dose-dependent manner. Since SSRIs have relatively long half-lives—often several days, and even longer for some of their active metabolites—and since it is well documented that SSRI concentrations in patients increase over time, it is assumed that the SSRI-TRKB interaction occurs only after the SSRI has accumulated sufficiently within the brain. As Casarotto *et al.* (2021) noted in the seminal article describing the discovery of the SSRI-TRKB interaction: “*the clinical response is only achieved when the drug reaches a brain concentration high enough to interact with a low-affinity binding target, such as TRKB*”.

While the data presented by Casarotto *et al.* demonstrate that SSRIs can interact with TRKB receptors under specific experimental conditions, our analyses—incorporating clinical measurements of brain SSRI concentrations, assessments of NMR sensitivity and spatial limits, reviews of pharmacodynamic data from rodent-derived neuronal *in vitro* studies, and our here presented own electrophysiological experiments in human iPSC-derived neurons—indicate that such an SSRI-TRKB interaction is highly unlikely to occur in human treated with SSRIs at clinically relevant doses.

## A Clinical-Experimental Gap: Therapeutic SSRI extracellular brain Levels Do Not Reach Micromolar Ranges

The clinical relevance of the SSRI-TRKB-receptor interaction has been inferred from fluoxetine-based magnetic resonance spectroscopy (MRS) studies, which estimated micromolar concentrations in the brains of chronically (multi-week) SSRI-treated individuals (Table 1, Komoroski *et al.*, 1994; Bolo *et al.*, 2000; Henry *et al.*, 2005).

**Table 1.** Reported concentrations of selective serotonin reuptake inhibitors (SSRIs) measured in human cerebrospinal fluid (CSF) or brain tissue. Concentrations (nM) are presented as mean ± standard deviation, as reported in the original studies.

| Bio sample type | Compound | Dose mg/day | Time of sampling after medication | Concentration nM | Reference |
| --- | --- | --- | --- | --- | --- |
| CSF | Citalopram | 20<br>20, 40 from day 5 | 4 weeks | 110 ± 71.0<br>91 ± 27 | (Paulzen et al., 2016)<br>(Nikisch, Eap and Baumann, 2008) |
| CSF | Paroxetine | 20, 30 from day 5 | 3 weeks | 5.590 ± 3.666 | (Lundmark et al., 1994) |
| CSF | Fluoxetine | 40 | 6 weeks | 26.5 ± 12.5 | (Mårtensson et al., 1989) |
| CSF | Norfluoxetine | 40 | 6 weeks | 16.9 ± 11.5 | (Mårtensson et al., 1989) |
| Brain ($^{19}\text{F}$ MRS) | Fluoxetine | 20 or 40<br>10 to 40<br>40 | 1 to 118 weeks<br>1 to 12 months<br>19 weeks | 9079.626 ± 4895.902<br>1300 ± 70<br>15655.25 ± 7192.478<br>13318.55 ± 11208.13 | (Karson et al., 1993)<br>(Bolo et al., 2000)<br>(Komoroski et al., 1994)<br>(Johnson, Lewis and Angier, 2007) |

Fluoxetine-specific MRS measurements date back to the 1990s and were pioneered by the team led by Komoroski. Still today, the technique is conducted using an RF head coil to excite and detect the proton resonances of fluoxetine within a localized brain voxel. The procedure begins with phantom measurements containing known concentrations of fluoxetine, which are scanned with the same coil and pulse sequence to identify its characteristic spectral peaks and to establish calibration curves.

During *in vivo* acquisition, the subject is positioned in the scanner, the magnetic field is shimmed to optimize spectral linewidth, and a PRESS or STEAM sequence is applied to isolate the region of interest; the resulting spectra are quantified by comparison to the phantom data. In the 1990s, the spatial resolution of these measurements was limited to relatively large voxels (≈8–20 cm³) due to low field strengths and early coil and analysis methods (Komoroski *et al.*, 1991). Today, improvements in magnet strength (3T and 7T), coil design, and spectral-fitting algorithms allow voxel sizes of 1–8 cm³ at 3T and ∼0.5–1.5 cm³ at 7T. In clinical settings, however, the typical voxel size remains around 2–8 cm³, balancing spatial resolution with the signal-to-noise required for quantification. Since the provide signals represent the aggregate magnetic resonance properties of large tissue volumes, this method cannot distinguish whether the detected molecules, e.g. Fluoxetine, are located within the interstitial space around neurons or glia, or in the blood within the cerebral vasculature (Lee, Adany and Choi, 2017; Maxouri *et al.*, 2023; Malaquin *et al.*, 2024). Beyond its spatial limitations, the method has low sensitivity and requires micromolar to millimolar reference solutions. Despite the absence of endogenous background signal, in vivo ¹⁹F-MRS cannot detect fluoxetine at micro- or nanomolar levels, as its signal remains far below the noise floor of current 1.5T–7T systems; only high-micromolar to low-millimolar concentrations are detectable. Thus, this imaging approach has substantial limitations in determining accurate drug concentrations. Moreover, factors such as magnetic field inhomogeneity, partial volume effects, local tissue composition, and the low signal-to-noise ratio for fluorine-based compounds further compromise quantification accuracy (Modo, 2021; Maxouri *et al.*, 2023; Du *et al.*, 2025). Consequently, these methods provide only an indirect and approximate assessment of brain SSRI levels (Karson *et al.*, 1993; Doyle *et al.*, 1995; Lee, Adany and Choi, 2017).

Extensive data are available from blood samples, which consistently show SSRI concentrations in the nanomolar range, with rare cases approaching 900 nM (Table 2) in chronic SSRI treated patients.Given the high plasma protein binding of SSRIs (e.g., more than 90% to albumin and other proteins) (Food and Drug Administration, 2023; van Harten, 1993; Vaswani, Linda and Ramesh, 2003), this is further argument of that is unlikely that neurons in the human brain are exposed to micromolar concentrations of SSRI, even at peak plasma levels (Table 2, Food and Drug Administration, 2023).

**Table 2.** Reported plasma concentrations of selective serotonin reuptake inhibitors (SSRIs) in humans under steady-state therapeutic dosing. Concentrations (nM) are presented as ranges or mean ± standard deviation, as reported in the original studies.

| SSRI | Dose mg/day | Time of sampling after medication | Concentration nM | Reference |
| --- | --- | --- | --- | --- |
| Citalopram | 20, 40 from day 5<br>20 | 4 weeks | 190 ± 629<br>348 ± 217 | (Nikisch, Eap and Baumann, 2008)<br>(Paulzen et al., 2016) |
| Paroxetine | 20, 30 from day 5 | 3 weeks | 236 ± 101 | (Lundmark et al., 1994) |
|  | 20, 40 after the first week | 6 weeks | 190 ± 114 | (Yasui-Furukori et al., 2011) |
| Fluoxetine | 20 | 8 weeks | 281 ± 147 | (Jd et al., 1997) |
|  | 20 or 40 | 1 to 118 weeks | 642 ± 217 | (Karson et al., 1993) |
|  | 10 to 40 | 1 to 12 months | 1730 ± 1000 | (Bolo et al., 2000) |
|  | 20 | 14 weeks | 266 ± 163 | (Brunswick et al., 2002) |
|  | 20 | 36 to 37 weeks | 152 ± 107 | (Heikkinen et al., 2003) |
|  | 40 | 6 weeks | 854 ± 359 | (Mårtensson et al., 1989) |
|  | 40 | 6 to 8 hours | 43 to 159 | (Food and Drug Administration, 2023) |
| Norfluoxetine | 40 | 6 weeks | 1006 ± 381 | (Mårtensson et al., 1989) |
|  | 20 | 36 to 37 weeks | 364 ± 73 | (Heikkinen et al., 2003) |

Although human CSF sampling studies are limited, it arguably represents the best approach to assess the extracellular concentration of SSRIs within the human brain. The rationale is that the interstitial fluid surrounding brain cells exchanges directly with the CSF, allowing CSF measurements to more accurately reflect extracellular SSRI levels (Margetis and Baker, 2025). Human CSF studies show that fluoxetine and other SSRIs are present only at pico-to nanomolar concentrations after several weeks of treatment (Table 1).

Here, we also discuss the theoretical scenario that SSRIs accumulate within the neuronal cell membrane and thereby reach sufficient concentrations to interact directly with the TRKB receptor. But do SSRIs bind and accumulate at the cell surface, within the membrane, inside the cell, or do they remain only in the extracellular space? While it has been demonstrated that SSRIs such as fluoxetine and escitalopram can permeate the cell membrane and transiently associate with lipid membranes when present extracellularly, recent evidence indicates that this accumulation is not sustained. In detail, Nichols *et al.* studied escitalopram and fluoxetine using newly developed, intensity-based, drug-sensing fluorescent reporters targeted to the plasma membrane, cytoplasm, and endoplasmic reticulum (ER) in cultured neurons and mammalian cells. Their data, supported by chemical detection of intracellular drug levels, showed that both SSRIs rapidly reach equilibrium between the extracellular solution and the neuronal cytoplasm or ER, with time constants of a few seconds for escitalopram and 200-300 seconds for fluoxetine. Although both drugs transiently accumulate in lipid membranes (approximately 18-fold for escitalopram and 180-fold for fluoxetine), they also leave the cytoplasm, organelles, and membranes just as rapidly upon washout (Nichols *et al.*, 2023). Thus, SSRIs do not persistently accumulate in the neuronal plasma membrane or within neurons under physiological conditions, indicating that they primarily accumulate within the extracellular space. To summarize, therapeutic SSRI extracellular brain levels do not reach micromolar ranges.

## Supratherapeutic Dosing in Rodent Studies Undermines SSRI-TRKB Translational Claims

The translational validity of proposed SSRI-TRKB interactions based on rodent behavioral studies warrants careful reconsideration. A recent review by Anderson *et al*. (2025) of 202 fluoxetine studies in rodents revealed that the median dose used (10 mg/kg) exceeded the animal-equivalent human dose by 3.2–6.5-fold, based on standard allometric scaling. This pattern was further supported by two systematic comparisons of dosing ranges across commonly used behavioral assays and antidepressant paradigms. In both cases, rodent fluoxetine exposures were found to be substantially higher than those achieved in humans.

Pharmacokinetic data underscore this discrepancy. An acute 20 mg oral dose of fluoxetine in humans yields plasma levels of 11.7–19.1 ng/ml, whereas an acute 10 mg/kg oral dose in rats results in 61.9 ng/ml. Under chronic dosing conditions, the widely used 10 mg/kg/day i.p. regimen in mice produces plasma concentrations more than an order of magnitude higher (1835.5 ng/ml) than those observed in humans receiving 20 mg/day (88.8–112.6 ng/ml). Compounding this issue, the majority of preclinical studies did not provide a rationale for the selected doses, and when a justification was given, it typically referenced prior literature rather than pharmacokinetic principles.

These considerations are particularly relevant for interpretations of SSRI-TRKB interactions. Casarotto *et al.* employed 15 mg/kg fluoxetine over two to three weeks to demonstrate behavioral effects dependent on TRKB. However, when scaled to human dosing, this regimen corresponds to supratherapeutic fluoxetine exposures unlikely to be achieved in the human brain. Consequently, while such studies may demonstrate pharmacological plausibility under high drug concentrations, they provide limited evidence for a mechanism operating within the concentration range relevant to human SSRI treatment.

Together, these findings highlight a broader challenge in the field: the reliance on rodent dosing paradigms that produce plasma and brain drug levels far exceeding those attainable in patients. This discrepancy severely limits the translational relevance of preclinical evidence supporting SSRI-TRKB interactions as a mechanism underlying antidepressant action in humans.

## Off-Target Actions of Micromolar SSRIs on Neuronal Ion Channels

Even in a theoretical scenario where human neurons are exposed to micromolar concentrations of SSRIs such as fluoxetine, it is highly likely that such elevated levels would exert adverse effects on neuronal function. Several studies have investigated how SSRIs influence neuronal activity using cell lines overexpressing specific voltage-gated ion channel subtypes, as well as primary rodent neuronal cultures, employing methods such as patch-clamp recordings and microelectrode arrays (Table 3). These *in vitro* electrophysiological approaches have revealed several off-target effects of SSRIs on neuronal activity, including voltage-gated sodium channel-mediated impairment of action potential generation and voltage-gated calcium channel-dependent modulation of neuronal excitability. All reported adverse or off-target effects of SSRIs occur at micromolar concentrations (Hahn *et al.*, 1999; Deák *et al.*, 2000; Igelström and Heyward, 2012; Poulin *et al.*, 2014). In overexpression systems, however, assays are typically conducted at high micromolar levels (10–100 µM) or higher. It is well established that overexpression of VGCCs in heterologous systems such as HEK293 cells necessitates 10-to 20-fold higher drug concentrations compared with assays performed in primary neurons. This discrepancy arises because numerous VGCC-interacting proteins present in neurons—but absent in cell lines—modulate channel kinetics and pharmacological sensitivity (Taglialatela, 2003; Kurejová *et al.*, 2007; Archana, Arunkumar and Omkumar, 2022).

**Table 3.** Reported neuronal ion channel targets of fluoxetine and other selective serotonin reuptake inhibitors (SSRIs). For each compound, the table lists the identified ion channels, the observed pharmacological effect, the effective concentration (µM), presented as IC_50_/EC_50_ values, and the experimental method employed, as described in original studies.

| SSRI | Target | Effect | Concentration μM | Method | Reference |
| --- | --- | --- | --- | --- | --- |
| Fluoxetine | Voltage-gated K<sup>+</sup> currents | Inhibition → increase in excitability | IC<sub>50</sub> = 16 | Whole-cell patch clamp | (Hahn et al., 1999) |
| Fluoxetine | Kv1.1 | Inhibition → increase in excitability | IC<sub>50</sub> > 500 | Whole-cell voltage clamp on oocytes | (Tytgat, Maertens and Daenens, 1997) |
| Fluoxetine | K<sub>v</sub>1.3 | Inhibition → increase in excitability | IC<sub>50</sub> = 5.9 | whole-cell voltage-clamp on CHO cells expressing K<sub>v</sub>1.3 | (Choi et al., 1999) |
| Fluoxetine | Kv1.4 | Inhibition → increase in excitability | IC<sub>50</sub> = 33.1 | Whole-cell patch clamp on CHO cells expressing K<sub>v</sub>1.4 | (Choi et al., 2003) |
| Fluoxetine | Kv1.5 | Inhibition → increase in excitability | IC<sub>50</sub> = 21 | Whole-cell patch clamp on CHO cells expressing K<sub>v</sub>1.5 | (Perchenet et al., 2001) |
| Fluoxetine | K<sub>v</sub>2.1 | Inhibition → increase in excitability | IC<sub>50</sub> = 51.3 | Whole-cell patch clamp on HEK293 cells expressing K<sub>v</sub>2.1 | (Chen et al., 2015; Wang et al., 2020) |
| Sertraline | K<sub>v</sub>2.1 | Inhibition → increase in excitability | IC<sub>50</sub> = 0.5 | Whole-cell and inside-out patch clamp on HEK293 cells expressing K<sub>v</sub>2.1 | (Delgado-Ramírez, López-Serrano and Rodríguez-Menchaca, 2024) |
| Fluoxetine | K<sub>v</sub>3.1 | Inhibition → increase in excitability | IC<sub>50</sub> = 13.11 | Whole-cell patch clamp on CHO cells expressing K<sub>v</sub>3.1 | (Choi et al., 2001) |
| Fluoxetine | K<sub>v</sub>4.3 | Inhibition → increase in excitability | IC<sub>50</sub> = 11.8 | Whole-cell patch clamp on CHO cells expressing K<sub>v</sub>4.3 | (Jeong, Choi and Hahn, 2013) |
| Fluoxetine | A-type K<sup>+</sup> currents | Inhibition → increase in excitability | IC<sub>50</sub> = 5.54 | Whole-cell patch clamp on rat brain slices | (Choi et al., 2004) |
| Fluoxetine | GIRK1/2 receptors (activated by 5-HT1A receptors) | Inhibition → increase in excitability | IC<sub>50</sub> = 16.9 | Whole-cell voltage clamp on oocytes expressing GIRK1/2 | (Kobayashi, Washiyama and Ikeda, 2003) |
| Fluoxetine | GIRK2 | Inhibition → increase in excitability | IC<sub>50</sub> = 89.5 | Whole-cell voltage clamp on oocytes expressing GIRK2 | (Kobayashi, Washiyama and Ikeda, 2003) |
| Citalopram | GIRK1/2 and GIRK1/4 | Inhibition → increase in excitability | 10 – 100 | Whole-cell voltage clamp on oocytes expressing GIRK1/2 | (Kobayashi, Washiyama and Ikeda, 2004) |
| Fluoxetine | Voltage-gated Na<sup>+</sup> currents | Inhibition → decrease in excitability | IC<sub>50</sub> = 25.6 | Whole-cell patch clamp | (Hahn et al., 1999) |
| Fluoxetine | Na<sub>v</sub>1.5 | Inhibition → decrease in excitability | IC<sub>50</sub> = 40 (-140 mV)<br>IC<sub>50</sub> = 4.7 (-90 mV) | Whole-cell patch clamp on HEK293 cells expressing Na<sub>v</sub>1.5 | (Poulin et al., 2014) |
| Fluoxetine | I<sub>NaP</sub> | Inhibition → decrease in excitability | At 10 | Electrophysiological recordings on alive brain slices | (Igelström and Heyward, 2012) |
| Fluoxetine | Voltage-gated $\text{Ca}^{2+}$ currents | Inhibition → decrease in excitability | $\text{IC}_{50} = 13.4$ | Whole-cell patch clamp | (Hahn et al., 1999) |
| Fluoxetine | T-type $\text{Ca}^{2+}$ currents | Inhibition → decrease in excitability | $\text{IC}_{50} = 6.8$ | Whole-cell patch clamp on rat hippocampal and prefrontal cells | (Deák et al., 2000) |
| Fluoxetine | HVA $\text{Ca}^{2+}$ currents, both L- and N-type | Inhibition → decrease in excitability | $\text{IC}_{50} = 1-2$ | Whole-cell patch clamp on rat hippocampal and prefrontal cells | (Deák et al., 2000) |
| Fluoxetine | GABA<sub>A</sub> receptor | Increase of response to GABA | $\text{EC}_{50} = 134.3 \pm 35.1 \mu\text{M}$ | Whole-cell patch clamp on HEK293T cells expressing GABA<sub>A</sub> receptor | (Robinson, Drafts and Fisher, 2003) |
| Fluoxetine | TREK-1 | Inhibition → increase in excitability | $\text{IC}_{50} = 19 \mu\text{M}$ | Whole-cell patch clamp on HEK293 cells expressing TREK-1, western blot on brain tissue | (Kennard et al., 2005; Chen et al., 2015) |
| Paroxetine | TREK-1 | Inhibition → increase in excitability | $\text{IC}_{50} = 5.5 \mu\text{M}$ | Whole-cell voltage clamp on DRN 5-HT and CA3 pyramidal neurons | (Heurteaux et al., 2006) |
| Sertraline | TREK-1 | Inhibition → increase in excitability | $\text{IC}_{50} = 3.2 \mu\text{M}$ | Whole-cell voltage clamp on DRN 5-HT and CA3 pyramidal neurons | (Heurteaux et al., 2006) |
| Fluoxetine | TREK-2 | Inhibition → increase in excitability | $\text{IC}_{50} = 28.7 \pm 7.6 \mu\text{M}$ | Whole-cell patch clamp on HEK293 cells expressing TREK-2 | (Kim et al., 2017) |
| Fluoxetine | nACh receptors | Inhibition → decrease in excitability | $\text{IC}_{50} = 1.6 \mu\text{M}$<br>$\text{IC}_{50} = 2-10 \mu\text{M}$ | Single-channel recordings on oocytes expressing nAChRs using the cell-attached mode<br>Whole-cell voltage clamp on oocytes expressing nAChRs | (Maggi et al., 1998)<br>(García-Colunga, Awad and Miledi, 1997) |

Not all SSRI-induced effects on specific VGCC subtypes have been independently replicated in primary neuronal cultures. Overall, SSRIs influence multiple VGCCs—core regulators of neuronal excitability and synaptic function—at micromolar, not nanomolar, concentrations. Furthermore, for certain reported effects observed at high micromolar concentrations (>10 µM) in heterologous expression systems, the corresponding effects in neurons are likely to occur at lower micromolar concentrations (1–10 µM) (Deák *et al.*, 2000; Kim *et al.*, 2013).

Two independent studies used primary cortical neurons cultured on microelectrode arrays to assess how fluoxetine impacts the neuronal network function of rodent neurons *in vitro* (Novellino *et al.*, 2011; Scelfo *et al.*, 2012). Importantly, the work described by Novellino *et al.* reported results from five independent laboratories (in Europe and the USA) that evaluated the impact of fluoxetine on the electrophysiological network function of primary rodent neuronal cultures.

A key advantage of this experimental approach is that it does not focus on a single VGCC subtype or neurotransmitter system. Instead, it captures the integrated activity of multiple VGCCs and both glutamatergic and GABAergic neurotransmitter systems present in neuronal cultures, thus preserving the physiological protein interactions that modulate VGCC and neurotransmitter receptor kinetics. In principle, this allows researchers to examine how SSRIs affect VGCCs and excitatory/inhibitory neurotransmission by detecting non-invasive changes in neuronal firing and network dynamics across multiple extracellular recording electrodes. Both studies—conducted across six independent laboratories—revealed that fluoxetine impairs neuronal network activity, specifically by reducing the mean firing rate and the number of network bursts, at micromolar but not nanomolar concentrations. While single-channel and overexpression studies provide valuable insight into how SSRIs act on individual molecular targets, they also reveal a major limitation: these studies isolate only one target at a time. As shown in Table 3, SSRIs interact with multiple VGCC subtypes, each producing distinct—and in some cases—opposing effects on neuronal excitability. For example, modulation of one channel subtype may enhance activity, whereas effects on another may suppress it. Consequently, single-channel assays cannot predict how these diverse interactions influence overall network activity.

Given this limitation, the use of microelectrode arrays (MEA) recordings becomes particularly important to assess the activity of neuronal networks. MEAs capture the activity of many interconnected neurons, allowing them to observe how multi-target drug interactions shape firing patterns and network dynamics. The interplay between multiple VGCC subtypes, neurotransmitter systems, and synaptic connections results in a neuronal population activity detectable by MEA technology. Thus, MEA recordings provide a more physiological meaningful assessment of how SSRIs impact neuronal network activity.

## MEA recordings combined with hiPSC-derived neurons reveal neuronal network dysfunction and inactivity at micromolar levels of SSRIs

Additionally, one could argue that these findings are derived from cell lines and animal in vitro neurons have limited translational value because human neurons may respond to SSRIs differently. A valid point as the impact of clinically relevant nanomolar SSRI concentrations on human neuronal function remains uncharacterized, particularly in comparison to the micromolar concentrations that have been shown to impair neuronal function in animal models but also needed for SSRI-TRKB interaction.

To address this translational gap, we employed human induced pluripotent stem cell (iPSC)-derived neurons cultured on microelectrode arrays (MEAs) to assess both single-neuron and network-level electrophysiological responses to nanomolar and micromolar SSRI concentrations. This approach models clinically relevant exposure levels as well as the higher concentrations associated with SSRI-TRKB binding and neuronal dysfunction.

The neural differentiation protocol used in this study generates telencephalic neural stem cells that give rise to cortical excitatory and inhibitory neurons, as characterized previously. Consequently, serotonin transporter (SERT) expression is absent, since cortical neurons do not express SERT and serotonergic presynaptic terminals are not present in this human in vitro model (Austin *et al.*, 1994; Borgers *et al.*, 2014). Nevertheless, this reductionist system is well suited to assessing how the electrophysiological function of human cortical excitatory and inhibitory neurons responds to nanomolar and micromolar SSRI exposure.

Three-dimensional (3D) human neural aggregates consisting of neurons, astrocytes, and oligodendroglial cells were generated from human induced pluripotent stem cells (iPSCs) and cultured on microelectrode arrays (MEAs) for four to five weeks, as previously described (Izsak *et al.*, 2019). The experiments were conducted on 40-day-old cultures, at which point the 3D human neural aggregates were fully developed. For network analysis, we evaluated activity detected simultaneously on the spatially separate electrodes embedded within each well of this setup. Network bursts, reflecting activity propagated through the neuronal network, were defined as periods of high-frequency spiking that occurred synchronously and are detected by spatial separated electrodes. Under standard culture condition, spontaneous activity revealed synchronous neuronal bursting across multiple electrodes, demonstrating the presence of a functionally connected neuronal network (Figure 1b).

**Figure 1.**
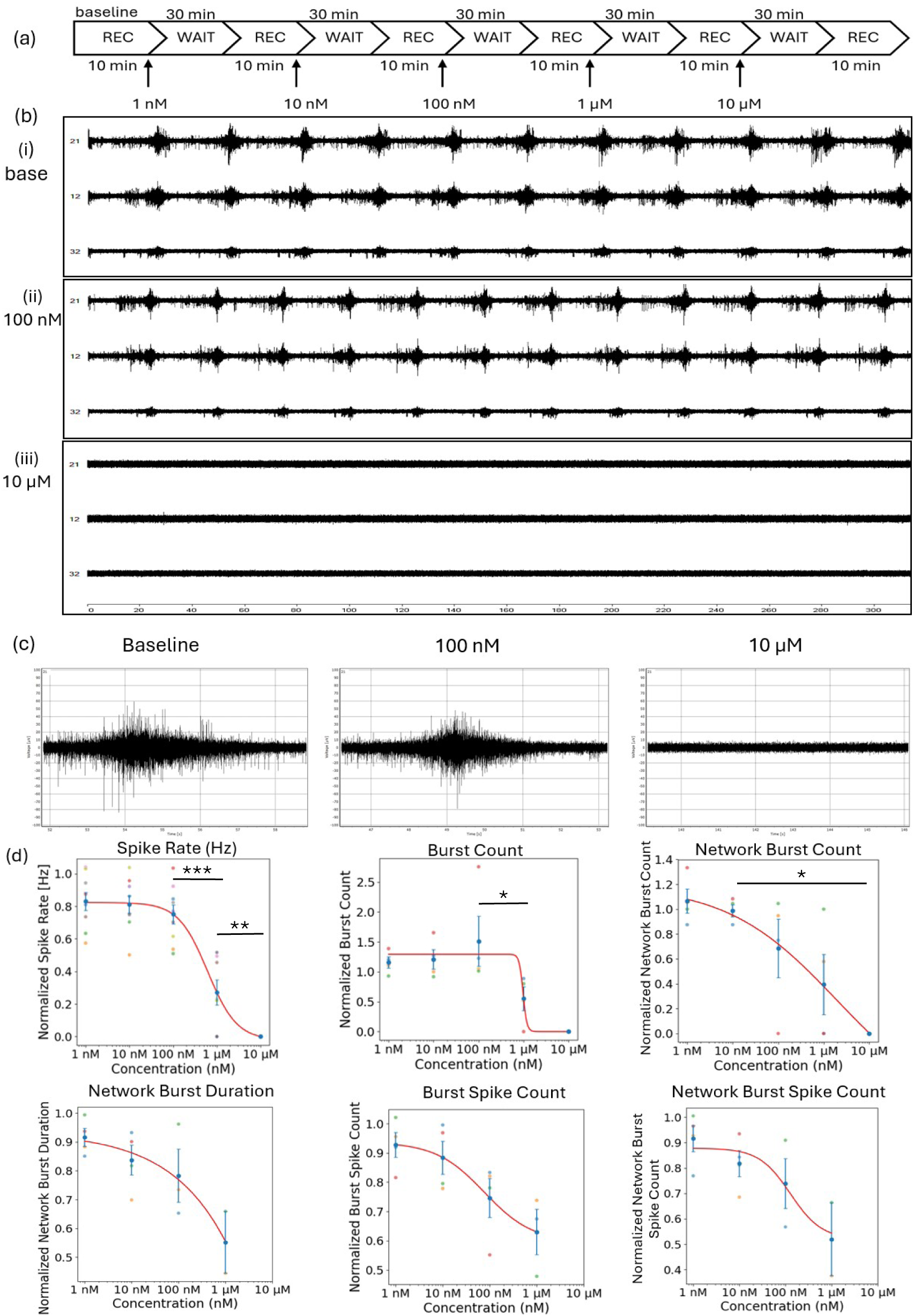
Effects of the SSRI fluoxetine hydrochloride on hiPSC-derived neuronal networks. (a) Schematic overview of the experimental workflow. (b) Representative microelectrode-array (MEA) recordings from three different electrodes obtained during (i) baseline, (ii) exposure to 100 nM fluoxetine, and (iii) exposure to 1 µM fluoxetine. (c) Representative network bursts detected during baseline, 100 µM fluoxetine, and 1 µM fluoxetine. (e) Dose-response relationships for spike rate (n = 9), burst count (n = 4), burst spike count (n = 4), network burst count (n = 4), network burst spike count (n = 4), and network burst duration (n = 4). All parameters are normalized to their respective baseline values. An error bar is presented as mean ± standard error of the mean. Statistical significance was evaluated using pairwise Mann-Whitney-U tests. p-values < 0.05 were considered significant (*), < 0.01 highly significant (**) and < 0.001 very highly significant (***) (Table S1).

This *in vitro* model, which has been extensively characterized in our laboratory, was employed for compound effect characterization. To assess the electrophysiological response of human neurons to selective serotonin reuptake inhibitors (SSRIs), we exposed cultures to therapeutically relevant nanomolar concentrations (1, 10, and 100 nM) as well as micromolar concentrations (1 and 10 µM) (Figure 1a).

Following the application of each concentration, cultures were incubated for 30 minutes prior to a 10-minute recording of neuronal network activity. Exposure to nanomolar concentrations of fluoxetine produced no detectable effects on neuronal electrophysiological properties. In contrast, exposure to 1 µM fluoxetine resulted in an approximate 75 % reduction in mean firing rate (Mann-Whitney U test, p = 0.000155, Table 4) and a 61 % reduction in the number of network bursts (Mann-Whitney U test, p = 0.066753, Table 4). Moreover, the mean duration of network bursts was reduced by 45 % (Mann-Whitney U test, p = 0.220671, Table 4) and the average number of spikes per network burst was reduced by 48 % (Mann-Whitney U test, p = 0.220671, Tab. 4) (Figure 1d).

**Table 4.** Statistical analysis of fluoxetine exposure in hiPSC-derived neuronal networks across six selected electrophysiological parameters: spike rate, burst count, burst spike count, network burst count, network burst duration, and network burst spike count. Statistical significance compared to baseline values was assessed using pairwise Mann-Whitney U tests. p-Values < 0.05 were considered statistically significant (*), whereas p-values > 0.05 were classified as non-significant (n. s.).

| Parameter | Concentration nM | p value | Significance |
| --- | --- | --- | --- |
| Spike Rate | 1 | 0.037949 | * |
|  | 10 | 0.003451 | ** |
|  | 100 | 0.003451 | ** |
|  | 1000 | 0.000155 | *** |
|  | 10000 | 0.000047 | *** |
| Burst Count | 1 | 0.281718 | n. s. |
|  | 10 | 0.619839 | n. s. |
|  | 100 | 0.021071 | * |
|  | 1000 | 0.021071 | * |
|  | 10000 | 0.013124 | * |
| Burst Spike Count | 1 | 0.281718 | n. s. |
|  | 10 | 0.021071 | n. s. |
|  | 100 | 0.021071 | * |
|  | 1000 | 0.063603 | * |
|  | 10000 |  | *** |
| Network Burst Count | 1 | 0.619839 | n. s. |
|  | 10 | 1.000000 | n. s. |
|  | 100 | 0.281718 | n. s. |
|  | 1000 | 0.066753 | n. s. |
|  | 10000 | 0.013124 | * |
| Network Burst Duration | 1 | 0.021071 | * |
|  | 10 | 0.021071 | * |
|  | 100 | 0.063603 | n. s. |
|  | 1000 | 0.220671 | n. s. |
|  | 10000 |  | *** |
| Network Burst Spike Count | 1 | 0.281718 | n. s. |
|  | 10 | 0.021071 | * |
|  | 100 | 0.063603 | n. s. |
|  | 1000 | 0.220671 | n. s. |
|  | 10000 |  | *** |

At 10 µM, fluoxetine exposure led to a complete abolition of neuronal network activity, with only occasional, uncorrelated spiking observed. Synchronous network bursting was entirely absent under these conditions (Figure 1b-c). Similar effects were observed for paroxetine (Figure S2, Figure S3).

To distinguish the effects of SSRIs on synaptic communication via neurotransmitter receptors and to specifically assess their impact on neuronal excitability under conditions in which neuronal activity is driven solely by VGCC function, we used a well-established experimental paradigm. In this paradigm, we applied blockers of fast glutamatergic and GABAergic transmission: CNQX and D-AP5 to block AMPA/kainate and NMDA receptor function, respectively, and picrotoxin to block GABA_A_ receptor function. These experiments were conducted under extracellular calcium-free conditions. The combination of these blockers, together with the absence of extracellular calcium, ensures that calcium-dependent neurotransmitter release—as well as the key neurotransmitter-mediated synaptic responses—is eliminated. Under such conditions, only uncorrelated spiking generated by intrinsically active neurons can be detected, as their activity is largely driven by VGCC channels (Illes *et al.*, 2014). In cultures treated with fluoxetine, we observed reductions in both spike rate and spike count (Figure 2e). Exposure to 1 µM fluoxetine resulted in a 43 % decrease in spike rate relative to the baseline (paired t-test, p = 0.734170, Table 5), whereas the spike count decreased by 2.3 % (paired t-test, p = 0.591541, Table 5). We also compared the adjusted media conditions (calcium-free and NMDA/AMPA-blocked) with standard media conditions. As expected, network bursts were completely absent under adjusted conditions (Figure 2b). Additionally, the total spike count decreased by 44.99 % (paired t-test, p = 0.208879, Table 6), and the mean spike rate was reduced by 72.5 % (paired t-test, p = 0.024954, Table 6).

**Figure 2.**
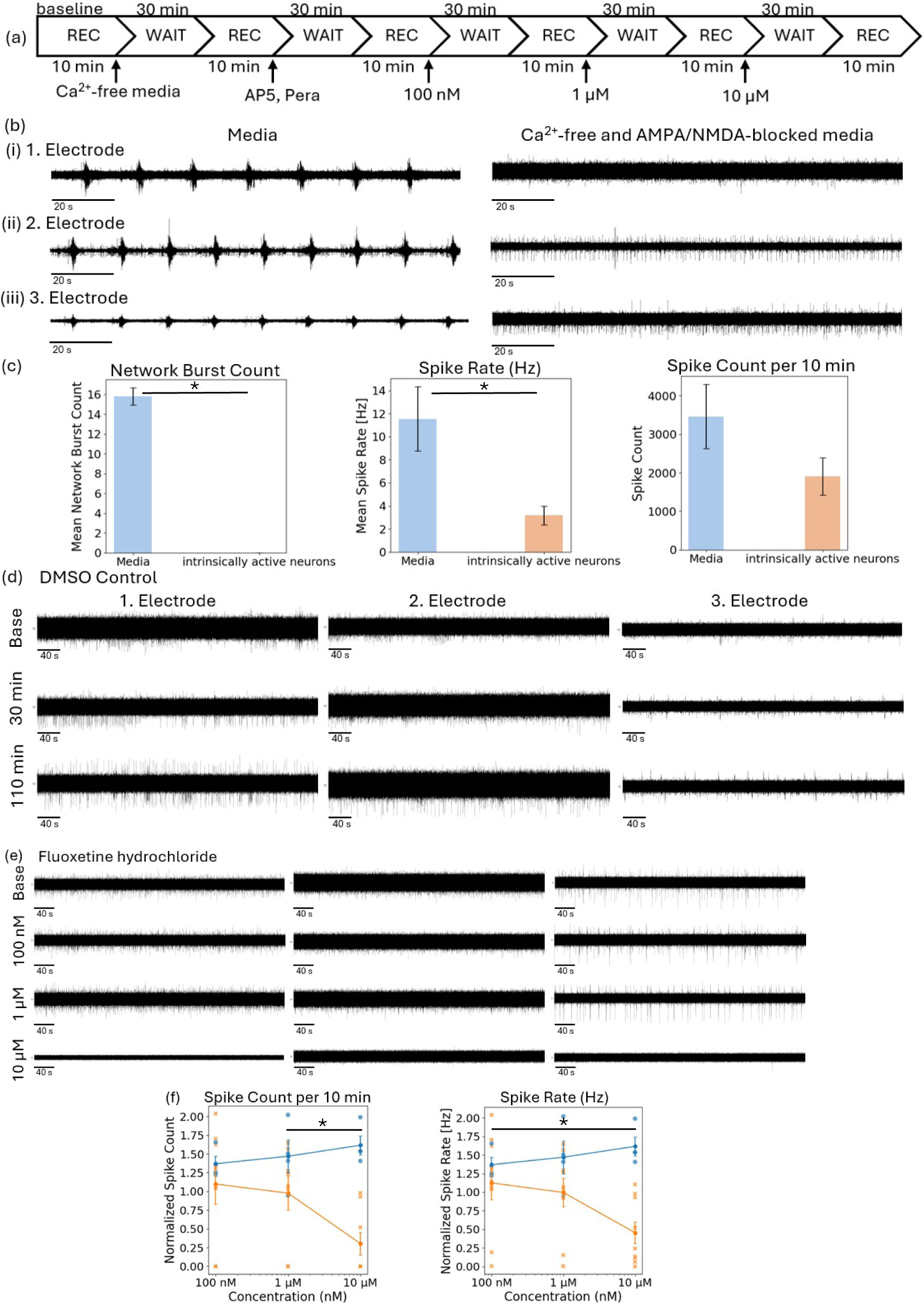
Effects of the SSRI fluoxetine hydrochloride on hiPSC-derived intrinsically active neurons. (a) Schematic overview of the experimental workflow. (bi-iii) Three representative MEA-recordings from electrodes belonging to three different MEA cultures under two conditions: standard media and Ca^2+^-free + NMDA/AMPA-blocked. (c) Bar graphs showing network burst count, spike count and spike rate for standard medium (blue, n = 10) and intrinsically active neurons (orange, n = 23). (d-e) Representative recordings from three selected electrodes under DMSO control (e) and fluoxetine treatment (e). (f) Spike count and spike rate for the DMSO control (blue, n = 4) and fluoxetine (orange, n = 8), represented as mean ± standard error of the mean. All parameters are normalized to their respective baseline values. Statistical significance was evaluated using pairwise Mann-Whitney-U tests. p-values < 0.05 were considered significant (*), < 0.01 highly significant (**) and < 0.001 very highly significant (***) (TabS3).

**Table 5.** Statistical analysis of fluoxetine exposure in hiPSC-derived intrinsically active neurons across three selected electrophysiological parameters: spike rate, spike count, and network burst count. Statistical significance compared to baseline values was assessed using paired t-tests. p-Values < 0.05 were considered significant (*), < 0.01 highly significant (**), and < 0.001 very highly significant (***). p-values > 0.05 were classified as non-significant (n. s.).

| Parameter | Concentration nM | p value | Significance |
| --- | --- | --- | --- |
| Spike Rate | 100 | 0.891416 | n. s. |
|  | 1000 | 0.734170 | n. s. |
|  | 10000 | 0.008683 | ** |
| Spike Count | 100 | 0.970410 | n. s. |
|  | 1000 | 0.591541 | n. s. |
|  | 10000 | 0.009117 | ** |

**Table 6.** Statistical analysis of hiPSC-derived intrinsically active neurons compared to standard media conditions across three selected electrophysiological parameters: spike rate, spike count, and network burst count. Statistical significance was assessed using paired t-tests. p-Values < 0.05 were considered significant (*), < 0.01 highly significant (**), and < 0.001 very highly significant (***). p-values > 0.05 were classified as non-significant (n. s.).

| Parameter | p value | Significance |
| --- | --- | --- |
| Spike Rate | 0.024954 | * |
| Spike Count | 0.208879 | n. s. |
| Network Burst Count | 0.000001475031 | *** |

## Are Therapeutic SSRI Concentrations Sufficient to Elicit SSRI-TRKB-Binding Effects on Patients?

This perspective does not seek to cast doubt on the discovery of the direct binding of SSRIs to the TRKB receptor or the resulting cellular and behavioral responses observed in animal models. The data presented by Casarotto *et al.* are robust and provide convincing evidence that SSRIs can directly interact with TRKB receptors.

However, what remains puzzling in clinical context is that this interaction requires micromolar concentrations of SSRIs, as unequivocally demonstrated by Casarotto *et al.* . When this is considered alongside the available evidence from clinical bio-sample analyses showing that extracellular brain SSRI concentrations in chronically treated patients remain well below the micromolar range, and with numerous reports documenting off-target effects of SSRIs on various neuronal ion channels at micromolar levels, the picture becomes complex.

Furthermore, the data presented here demonstrate that micromolar—but not clinically relevant nanomolar—SSRI concentrations abolish neuronal network activity in human iPSC-derived neurons. Therefore, based on the current evidence, we conclude that there is insufficient support for the notion that the SSRI-TRKB interaction occurs in the brains of patients treated with SSRIs.

## Supplementary Figures

**Figure S1.**
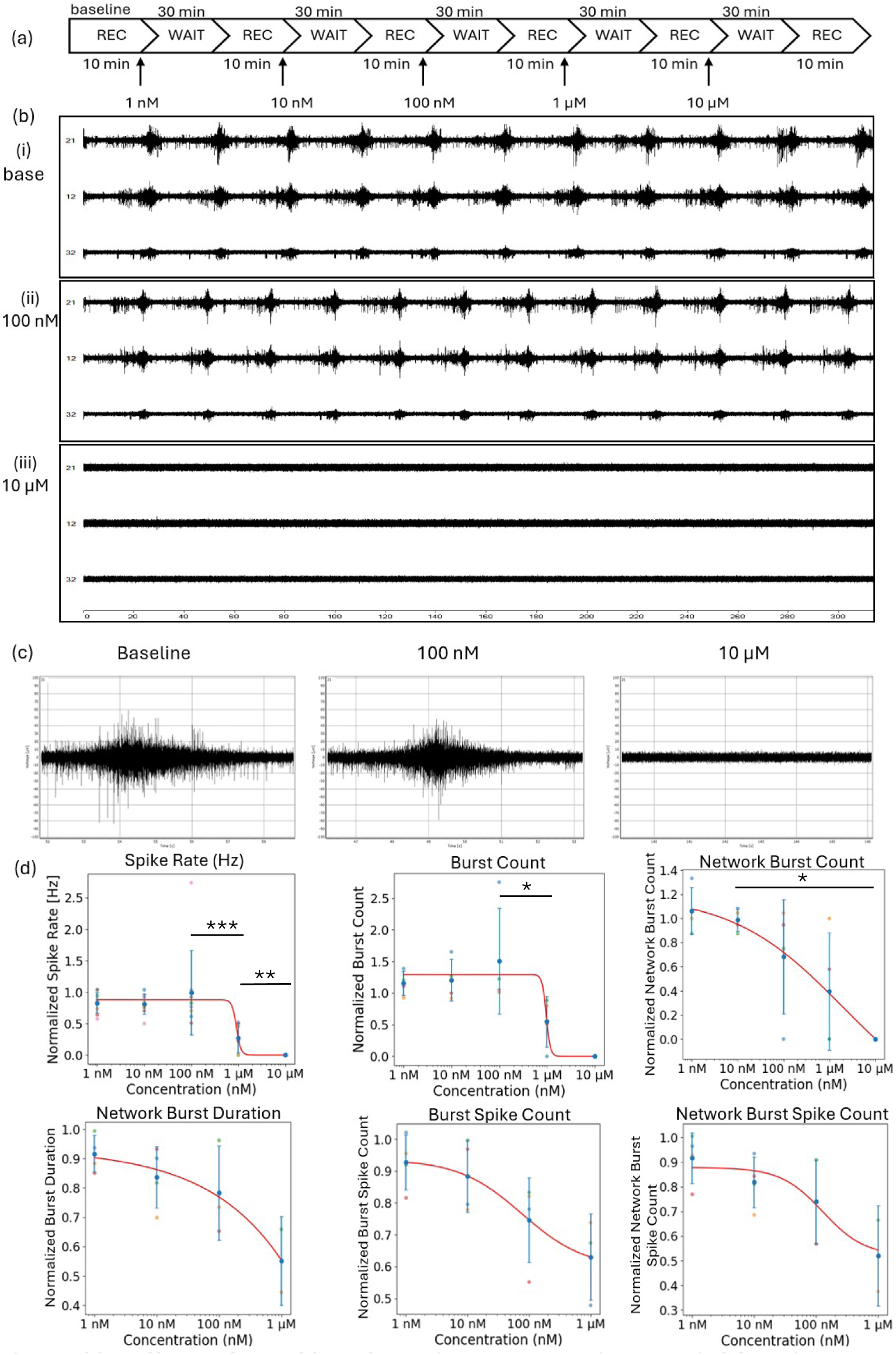
Effects of the SSRI fluoxetine hydrochloride on hiPSC-derived neuronal networks. (a) Schematic overview of the experimental workflow. (b) Representative microelectrode-array (MEA) recordings from three different electrodes obtained during (i) baseline, (ii) exposure to 100 nM fluoxetine, and (iii) exposure to 1 µM fluoxetine. (c) Representative network bursts detected during baseline, 100 µM fluoxetine, and 1 µM fluoxetine. (e) Dose-response relationships for spike rate (n = 9), burst count (n = 4), burst spike count (n = 4), network burst count (n = 4), network burst spike count (n = 4), and network burst duration (n = 4). All parameters are normalized to their respective baseline values. An error bar is presented as mean ± standard deviation. Statistical significance was evaluated using pairwise Mann-Whitney-U tests. p-values < 0.05 were considered significant (*), < 0.01 highly significant (**) and < 0.001 very highly significant (***).

**Figure S2.**
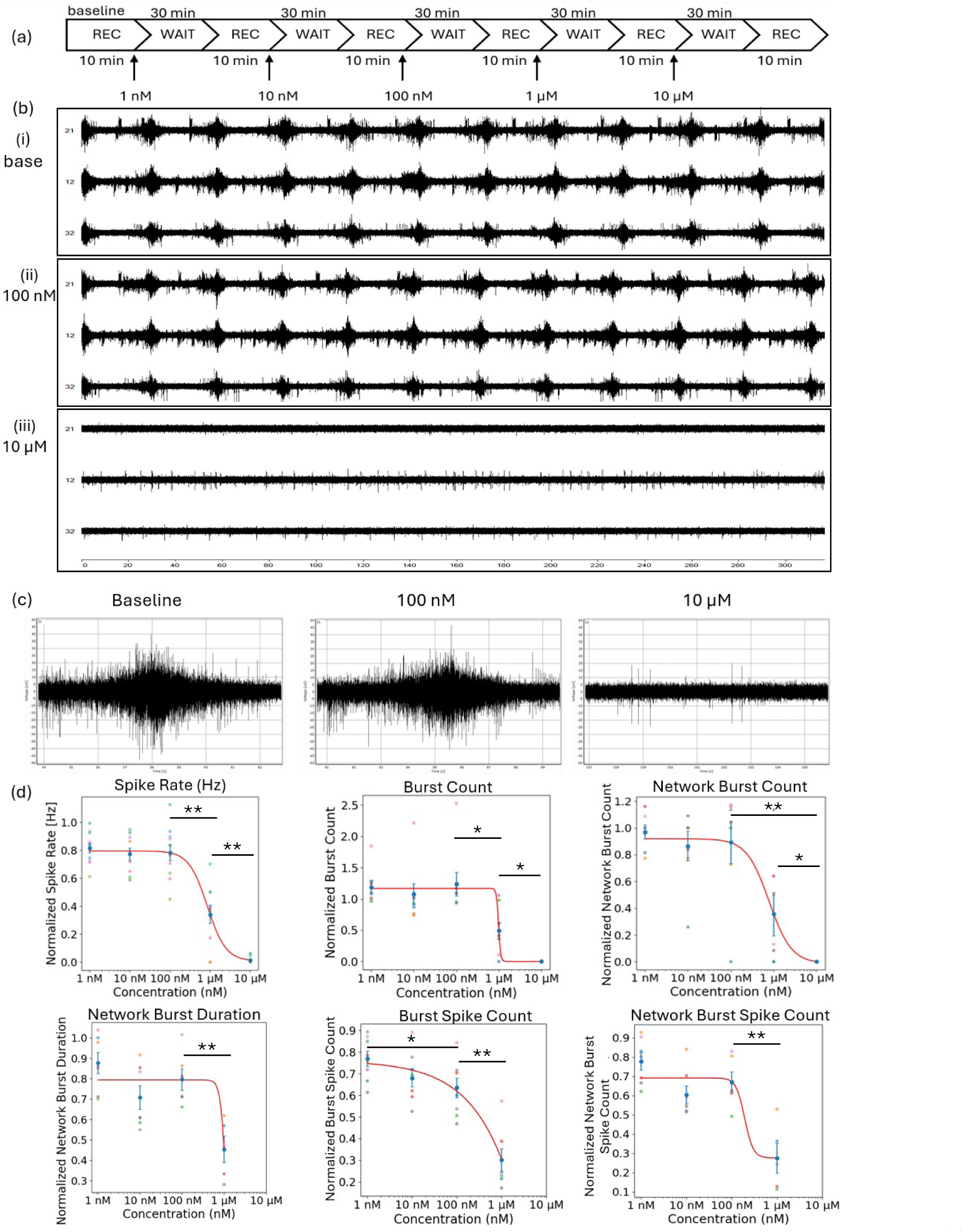
Effects of the SSRI paroxetine hydrochloride on hiPSC-derived neuronal networks. (a) Schematic overview of the experimental workflow. (b) Representative microelectrode-array (MEA) recordings from three different electrodes obtained during (i) baseline, (ii) exposure to 100 nM paroxetine, and (iii) exposure to 1 µM paroxetine. (c) Representative network bursts detected during baseline, 100 µM paroxetine, and 1 µM paroxetine. (e) Dose-response relationships for spike rate (n = 9), burst count (n = 4), burst spike count (n = 4), network burst count (n = 4), network burst spike count (n = 4), and network burst duration (n = 4). All parameters are normalized to their respective baseline values. An error bar is presented as mean ± standard error of the mean. Statistical significance was evaluated using pairwise Mann-Whitney-U tests. p-values < 0.05 were considered significant (*), < 0.01 highly significant (**) and < 0.001 very highly significant (***).

**Figure S3.**
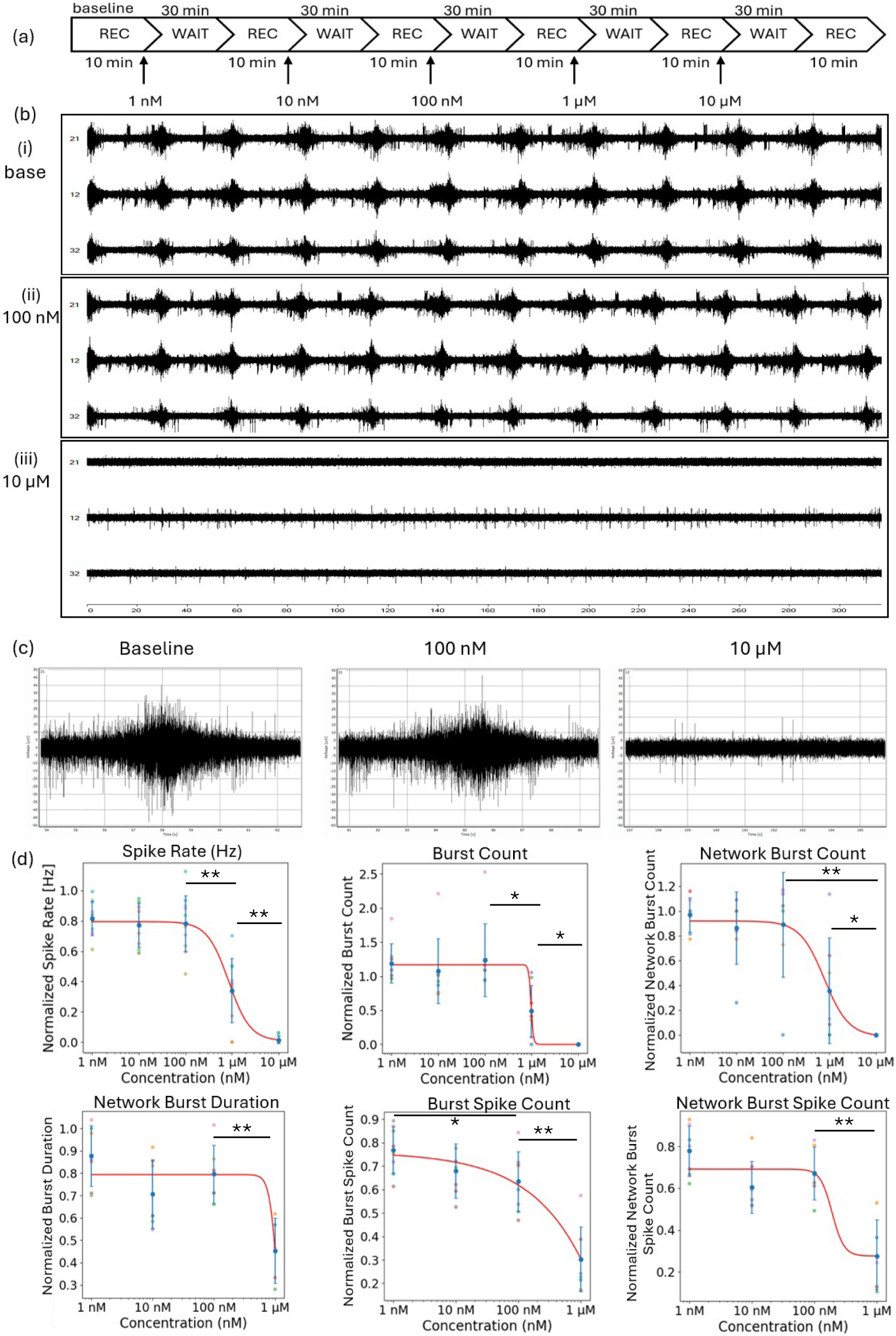
Effects of the SSRI paroxetine hydrochloride on hiPSC-derived neuronal networks. (a) Schematic overview of the experimental workflow. (b) Representative microelectrode-array (MEA) recordings from three different electrodes obtained during (i) baseline, (ii) exposure to 100 nM paroxetine, and (iii) exposure to 1 µM paroxetine. (c) Representative network bursts detected during baseline, 100 µM paroxetine, and 1 µM paroxetine. (e) Dose-response relationships for spike rate (n = 9), burst count (n = 4), burst spike count (n = 4), network burst count (n = 4), network burst spike count (n = 4), and network burst duration (n = 4). All parameters are normalized to their respective baseline values. An error bar is presented as mean ± standard deviation. Statistical significance was evaluated using pairwise Mann-Whitney-U tests. p-values < 0.05 were considered significant (*), < 0.01 highly significant (**) and < 0.001 very highly significant (***).

## Supplementary Tables

**Table S1.** Statistical analysis of fluoxetine and paroxetine exposure in hiPSC-derived neuronal networks across six selected electrophysiological parameters: spike rate, burst count, burst spike count, network burst count, network burst duration, and network burst spike count. Pairwise Mann-Whitney U tests were used to evaluate statistical significance. p-Values < 0.05 were considered significant (*), < 0.01 highly significant (**), and < 0.001 very highly significant (***). p-values > 0.05 were classified as non-significant (n. s.).

| Parameter | concentration pair | p value |  | Significance |  |
| --- | --- | --- | --- | --- | --- |
|  |  | Fluoxetine | Paroxetine | Fluoxetine | Paroxetine |
| Spike Rate | 1 nM vs 1 μM | 0.000401 | 0.000517 | *** | *** |
| | 10 μM vs 1 μM | 0.005317 | 0.002501 | ** | ** |
| | 10 μM vs 1 nM | 0.000161 | 0.000125 | *** | *** |
| | 10 nM vs 1 μM | 0.000773 | 0.000974 | *** | *** |
|  | 10 nM vs 1 nM | 0.859819 | 0.563618 | n. s. | n. s. |
| | 10 nM vs 10 μM | 0.000161 | 0.000125 | *** | *** |
| | 100 nM vs 1 μM | 0.000773 | 0.001194 | *** | ** |
|  | 100 nM vs 1 nM | 0.331387 | 0.644095 | n. s. | n. s. |
| | 100 nM vs 10 μM | 0.000161 | 0.000125 | *** | *** |
|  | 100 nM vs 10 nM | 0.536499 | 1.000000 | n. s. | n. s. |
| Burst Count | 1 nM vs 1 μM | 0.028571 | 0.001865 | * | ** |
| | 10 μM vs 1 μM | 0.068918 | 0.001460 | n. s. | * |
| | 10 μM vs 1 nM | 0.021071 | 0.000410 | * | *** |
| | 10 nM vs 1 μM | 0.028571 | 0.020668 | * | * |
|  | 10 nM vs 1 nM | 1.000000 | 0.114916 | n. s. | n. s. |
| | 10 nM vs 10 μM | 0.021071 | 0.000410 | * | *** |
| | 100 nM vs 1 μM | 0.028571 | 0.004662 | * | ** |
|  | 100 nM vs 1 nM | 0.885714 | 0.573737 | n. s. | n. s. |
| | 100 nM vs 10 μM | 0.021071 | 0.000410 | * | *** |
|  | 100 nM vs 10 nM | 0.685714 | 0.114916 | n. s. | n. s. |
| Burst Spike Count | 1 nM vs 1 μM | 0.057143 | 0.000311 | n. s. | *** |
| | 10 nM vs 1 μM | 0.057143 | 0.000622 | n. s. | *** |
|  | 10 nM vs 1 nM | 0.685714 | 0.160528 | n. s. | n. s. |
| | 100 nM vs 1 μM | 0.228571 | 0.002176 | n. s. | ** |
|  | 100 nM vs 1 nM | 0.114286 | 0.028127 | n. s. | * |
|  | 100 nM vs 10 nM | 0.485714 | 0.505361 | n. s. | n. s. |
| Network Burst Count | 1 nM vs 1 μM | 0.079594 | 0.044248 | n. s. | * |
| | 10 μM vs 1 μM | 0.185877 | 0.012777 | n. s. | * |
| | 10 μM vs 1 nM | 0.021071 | 0.001446 | * | ** |
| | 10 nM vs 1 μM | 0.110210 | 0.090247 | n. s. | n. s. |
|  | 10 nM vs 1 nM | 0.771503 | 0.832045 | n. s. | n. s. |
| | 10 nM vs 10 μM | 0.021071 | 0.001446 | * | ** |
| | 100 nM vs 1 μM | 0.459597 | 0.109550 | n. s. | n. s. |
|  | 100 nM vs 1 nM | 0.200000 | 0.751115 | n. s. | n. s. |
| | 100 nM vs 10 μM | 0.068918 | 0.004569 | n. s. | ** |
|  | 100 nM vs 10 nM | 0.306492 | 0.561211 | n. s. | n. s. |
| Network Burst Spike Count | 1 nM vs 1 μM | 0.133333 | 0.002525 | n. s. | ** |
| | 10 nM vs 1 μM | 0.133333 | 0.017677 | n. s. | * |
|  | 10 nM vs 1 nM | 0.342857 | 0.037879 | n. s. | * |
| | 100 nM vs 1 μM | 0.400000 | 0.008658 | n. s. | ** |
|  | 100 nM vs 1 nM | 0.114286 | 0.137529 | n. s. | n. s. |
|  | 100 nM vs 10 nM | 0.628571 | 0.445221 | n. s. | n. s. |
| Network Burst Duration | 1 nM vs 1 μM | 0.133333 | 0.002525 | n. s. | ** |
| | 10 nM vs 1 μM | 0.133333 | 0.047980 | n. s. | * |
|  | 10 nM vs 1 nM | 0.342857 | 0.053030 | n. s. | n. s. |
| | 100 nM vs 1 μM | 0.400000 | 0.004329 | n. s. | ** |
|  | 100 nM vs 1 nM | 0.400000 | 0.445221 | n. s. | n. s. |
|  | 100 nM vs 10 nM | 0.857143 | 0.294872 | n. s. | n. s. |

**Table S2.** Statistical analysis of fluoxetine- or paroxetine-treated and control hiPSC-derived neuronal networks across six selected electrophysiological parameters: spike rate, burst count, burst spike count, network burst count, network burst duration, and network burst spike count. Pairwise Mann-Whitney U tests were used to evaluate statistical significance. p-Values < 0.05 were considered significant (*), < 0.01 highly significant (**), and < 0.001 very highly significant (***). p-values > 0.05 were classified as non-significant (n. s.).

| Parameter | compound vs control pair | p value |  | Significance |  |
| --- | --- | --- | --- | --- | --- |
|  |  | Fluoxetine | Paroxetine | Fluoxetine | Paroxetine |
| Spike Rate | 1 nM | 0.376333 | 0.481499 | n. s. | n. s. |
|  | 10 nM | 0.455944 | 0.511857 | n. s. | n. s. |
|  | 100 nM | 0.516493 | 0.541835 | n. s. | n. s. |
| | 1 μM | 0.167538 | 0.236807 | n. s. | n. s. |
| | 10 μM | 0.008125 | 0.028973 | ** | ** |
| Burst Count | 1 nM | 0.057143 | 0.012121 | n. s. | * |
|  | 10 nM | 0.057143 | 0.084848 | n. s. | n. s. |
|  | 100 nM | 0.228571 | 0.193939 | n. s. | n. s. |
| | 1 μM | 1.000000 | 1.000000 | n. s. | n. s. |
| | 10 μM | 0.122745 | 0.023163 | n. s. | * |
| Burst Spike Count | 1 nM | 0.266667 | 0.888889 | n. s. | n. s. |
|  | 10 nM | 0.133333 | 0.711111 | n. s. | n. s. |
|  | 100 nM | 0.533333 | 0.533333 | n. s. | n. s. |
| | 1 μM | 1.000000 | 0.111111 | n. s. | n. s. |
| | 10 μM | | | *** | *** |
| Network Burst Count | 1 nM | 0.400000 | 0.845815 | n. s. | n. s. |
|  | 10 nM | 0.800000 | 1.000000 | n. s. | n. s. |
|  | 100 nM | 1.000000 | 0.845815 | n. s. | n. s. |
| | 1 μM | 0.716801 | 0.324723 | n. s. | n. s. |
| | 10 μM | 0.133614 | 0.013328 | n. s. | * |
| Network Burst Spike Count | 1 nM | 0.400000 | 0.250000 | n. s. | n. s. |
|  | 10 nM | 0.400000 | 0.250000 | n. s. | n. s. |
|  | 100 nM | 0.500000 | 0.285714 | n. s. | n. s. |
| | 1 μM | 1.000000 | 0.333333 | n. s. | n. s. |
| | 10 μM | | | *** | *** |
| Network Burt Duration | 1 nM | 0.400000 | 0.250000 | n. s. | n. s. |
|  | 10 nM | 0.400000 | 0.250000 | n. s. | n. s. |
|  | 100 nM | 0.500000 | 0.285714 | n. s. | n. s. |
| | 1 μM | 0.666667 | 0.333333 | n. s. | n. s. |
| | 10 μM | | | *** | *** |

**Table S3.**
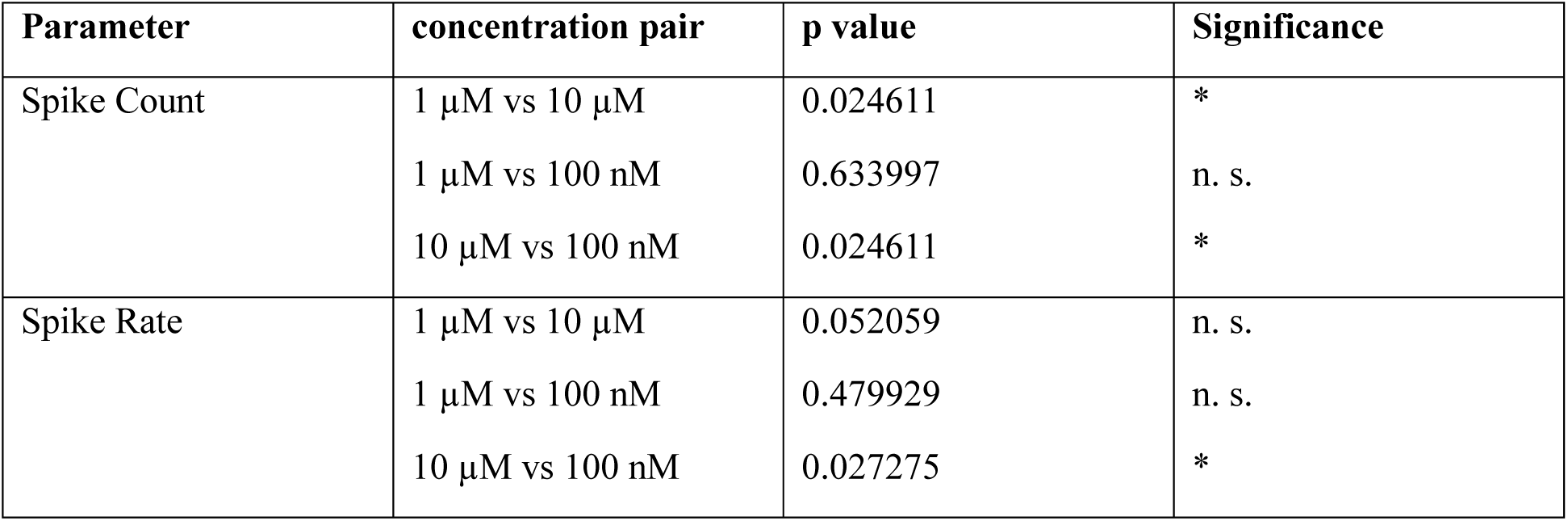
Statistical analysis of fluoxetine exposure in hiPSC-derived intrinsically active neurons across three selected electrophysiological parameters: spike rate, spike count, and network burst count. Pairwise Mann-Whitney U tests were used to evaluate statistical significance. p-Values < 0.05 were considered significant (*), whereas p-values > 0.05 were classified as non-significant (n. s.).

**Table S4.** Statistical analysis of fluoxetine-treated and control hiPSC-derived intrinsically active neurons across three selected electrophysiological parameters: spike rate, spike count, and network burst count. Pairwise Mann-Whitney U tests were used to evaluate statistical significance. p-Values < 0.05 were considered significant (*), < 0.01 highly significant (**), whereas p-values > 0.05 were classified as non-significant (n. s.).

| Parameter | compound vs control pair | p value | Significance |
| --- | --- | --- | --- |
| Spike Count | 100 nM | 0.670583 | n. s. |
| | 1 μM | 0.349394 | n. s. |
| | 10 μM | 0.006339 | ** |
| Spike Rate | 100 nM | 0.503497 | n. s. |
| | 1 μM | 0.260140 | n. s. |
| | 10 μM | 0.002797 | ** |

## Notes

### Competing Interest Statement

SI is the founder of Oscillution AB, holds shares in the company, and serves in a management position.

### Funding

- forska utan djurförsök, Sweden

